# Autoresearch Discovery of Interpretable Filter Rules for Antibody Binder Classification

**DOI:** 10.64898/2026.05.05.723069

**Authors:** Mikel Landajuela

**Affiliations:** Lawrence Livermore National Laboratory, USA

## Abstract

Antibody design campaigns increasingly generate many candidates before only a small subset can be tested experimentally, making candidate filtering a central bottleneck. We study whether an autoresearch loop can discover better training-free filters for antibody binder classification by iteratively proposing rule variants, evaluating them under a fixed Leave-One-System-Out protocol, recording each experiment in version control, and using the results to guide the next iteration. Across 75 unique logged filter variants on seven antibody-antigen systems, the loop improves average ROC-AUC from 0.6371 for the initial baseline to 0.8060 for a compact final rule that we call the *RMSD-Tuned Triad* rule, an absolute gain of 0.1689 and a relative improvement of 26.5%. The discovered filter is competitive with supervised machine learning baselines and prompted LLM baselines evaluated on the same systems: it exceeds logistic regression (0.7144), feature-selected balanced logistic regression (0.7536), and GPT-4o tabular few-shot prompting (0.7640), and it comes within 0.0044 ROC-AUC of the strongest GPT-5 tabular few-shot result (0.8104). Unlike the LLM baseline, the final rule requires no prompted examples and no LLM inference once the numeric structure-derived features are available. These results show that systematic autoresearch can turn simple structural-confidence signals into compact, interpretable filters that are useful when target-specific training data are scarce.

## 1 Introduction

Computational antibody design is often a generate-then-filter problem. Modern generators can propose large libraries of candidate complementarity-determining region (CDR) sequences, but only a small fraction can be synthesized and assayed. This pattern appears broadly in protein and antibody design: deep-learning-augmented binder design generates large computational libraries and uses structure-prediction filters to enrich experimental hit rates [1, 2]; tFold separates antibody generation from a downstream filter that uses confidence, structure, novelty, motif, and specificity criteria before wet-lab validation [3]; and constrained antibody-library optimization explicitly scores large mutational search spaces before selecting a smaller diverse library for testing [4]. Filtering is also common downstream in therapeutic antibody development, where computational and experimental developability screens flag liabilities such as aggregation, surface hydrophobicity, charge imbalance, expression risk, or poor manufacturability before costly scale-up [5, 6, 7].

This paper focuses on the filter itself. Rather than training a new binding model, we ask whether an autoresearch loop can discover a better rule-based classifier from existing structural and confidence features. The loop repeatedly modifies the filter, evaluates it on a fixed benchmark, records the result, and carries forward changes that improve cross-system ranking. The explored design choices include threshold calibration, continuous scoring in place of binary decisions, metric selection, non-linear score transformations, and metric weighting; these choices are evaluated as part of one iterative process rather than as a hand-picked final recipe.

We evaluate on seven systems from the IgDesign antibody design benchmark [8]. Each candidate has an experimental binder label. In addition to the benchmark metadata and antibody structure-ensemble RMSD metrics from the IgDesign workflow, we added AlphaFold3 complex-confidence features [9] and ProteinMPNN sequence-design scores [10] as part of this study. The main result is that a compact training-free filter discovered by autoresearch reaches 0.8060 average ROC-AUC, substantially improving over the initial rule baseline and nearly matching the strongest GPT-5 prompting baseline without using few-shot examples, binding labels for training, or LLM calls at inference time.

## 2 Autoresearch Loop

The autoresearch loop treats filter design as an empirical search problem, following the general pattern of agent-driven experiment proposal, execution, evaluation, and iteration popularized by Karpathy’s autoresearch repository [11]. Each iteration begins with a hypothesis about the scoring rule, such as replacing a hard cutoff with a continuous score or changing the relative weight of interface confidence and structural consistency. The implementation is committed with a descriptive message, evaluated with the same Leave-One-System-Out protocol, and appended to a results log containing the commit hash and summary metrics. The next experiment is chosen from the observed behavior of prior experiments, keeping the search interpretable and auditable.

The search space is intentionally small enough to inspect. A filter may use hard thresholds, continuous scores, or a hybrid of both; it may include AlphaFold3 interface confidence, global confidence, predicted aligned error, local confidence, ensemble RMSD, and sequence designability scores; and it may combine normalized signals additively, multiplicatively, or after power transformations. This design makes each experiment easy to explain, while still allowing the loop to find non-obvious combinations that outperform literature-style threshold rules.

## 3 Experimental Setup

### 3.1 Dataset and Features

#### 3.1.1 IgDesign Antibody-Antigen Systems

We evaluate on seven antibody-antigen systems from the IgDesign dataset. Each system contains variants of a reference antibody-antigen pair and binary experimental binding labels. The systems differ substantially in binder prevalence, which makes random sample-level cross-validation too optimistic; throughout the paper, a full antibody-antigen system is held out at a time.

#### 3.1.2 Features

For each variant, we use numeric features produced by the existing structure-prediction and sequence-scoring pipeline. The autoresearch loop does not learn feature embeddings or use labels to fit model parameters; it only searches over how raw or normalized feature values are transformed and combined.

### 3.2 Evaluation Protocol

We use Leave-One-System-Out (LOSO) cross-validation. In each fold, one antibody-antigen system is held out for testing and the remaining six systems are available only for methods that require training. Rule-based filters are applied directly to the held-out system without fitting. Machine learning baselines are trained on the six non-held-out systems and evaluated on the held-out system. The primary metric is ROC-AUC because antibody screening is primarily a ranking task: experimentalists can test the highest-scoring candidates under a chosen budget, and ROC-AUC does not depend on a fixed classification threshold.

### 3.3 Baselines

We compare the autoresearch filter with two conventional baseline groups. The first group contains adapted rule-based filters: the initial baseline, AF2-IG-easy, Protenix-style, AF3-style, and PXDesign-style thresholds. These filters follow the confidence- and geometry-threshold families summarized in the PXDesign filtering benchmark [12], where AF2-IG is traced to the AlphaFold2 initial-guess binder-design workflow [1], AF2-IG-easy follows BindCraft-style relaxed filtering [2], AF3-style filtering uses AlphaFold3 confidence metrics [9], and Protenix-style filters use Protenix confidence metrics [13]. The second group contains supervised logistic-regression baselines trained on LOSO training folds.

### 3.4 LLM Prompting Baselines

We also evaluate zero-shot LLM prompting baselines that consume the same candidate-level context used by the rule-based filters, augmented with compact sequence, mutation, and interface summaries. The **GPT-4o full** prompt includes verbose sequence context, contact graphs, interaction profiles, mutation lists, AlphaFold3 metrics, structure-ensemble summaries, and sequence-scoring summaries. The **GPT-4o features** prompt removes most narrative context and presents the same evidence as structured feature blocks. The **GPT-4o tabular+few-shot** prompt further compresses the input into numeric feature and delta tables and prepends two binder and two non-binder calibration examples from the same antigen system. The **GPT-5 tabular+few-shot** baseline uses the same tabular few-shot format with a larger model. For each LLM run, the model returns a binder probability; ROC-AUC is computed per held-out system from those probabilities and then averaged across the seven systems. No LLM baseline is fine-tuned on the binding labels.

## 4 Results

### 4.1 Performance Comparison

Table 3 reports the main comparison. The autoresearch filter improves over the initial baseline by +0.1689 ROC-AUC, or 26.5% relative to the 0.6371 baseline. It also exceeds the best adapted literature-style threshold baseline in this benchmark, Protenix-style filtering at 0.6612 ROC-AUC. Against learning-based comparators, it outperforms both supervised logistic-regression baselines and the GPT-4o LLM prompting variants, while remaining within 0.0044 ROC-AUC of GPT-5 tabular few-shot prompting. This makes the final rule practically comparable to the strongest LLM result, but with a much simpler deployment path: score the candidate table once, without few-shot calibration examples or LLM calls.

**Table 1.** Dataset statistics per antibody-antigen system.

| System | Samples | Binders | % Binders |
| --- | --- | --- | --- |
| Afasevikumab-IL17A | 179 | 6 | 3.4% |
| Bimagrumab-ACVR2B | 212 | 15 | 7.1% |
| Eculizumab-C5 | 198 | 30 | 15.2% |
| Osocimab-FXI | 116 | 34 | 29.3% |
| Spesolimab-IL36R | 177 | 29 | 16.4% |
| Tezepelumab-TSLP | 176 | 115 | 65.3% |
| Utomilumab-TNFRSF9 | 185 | 29 | 15.7% |
| **Total** | **1,243** | **258** | **20.8%** |

**Table 2.**
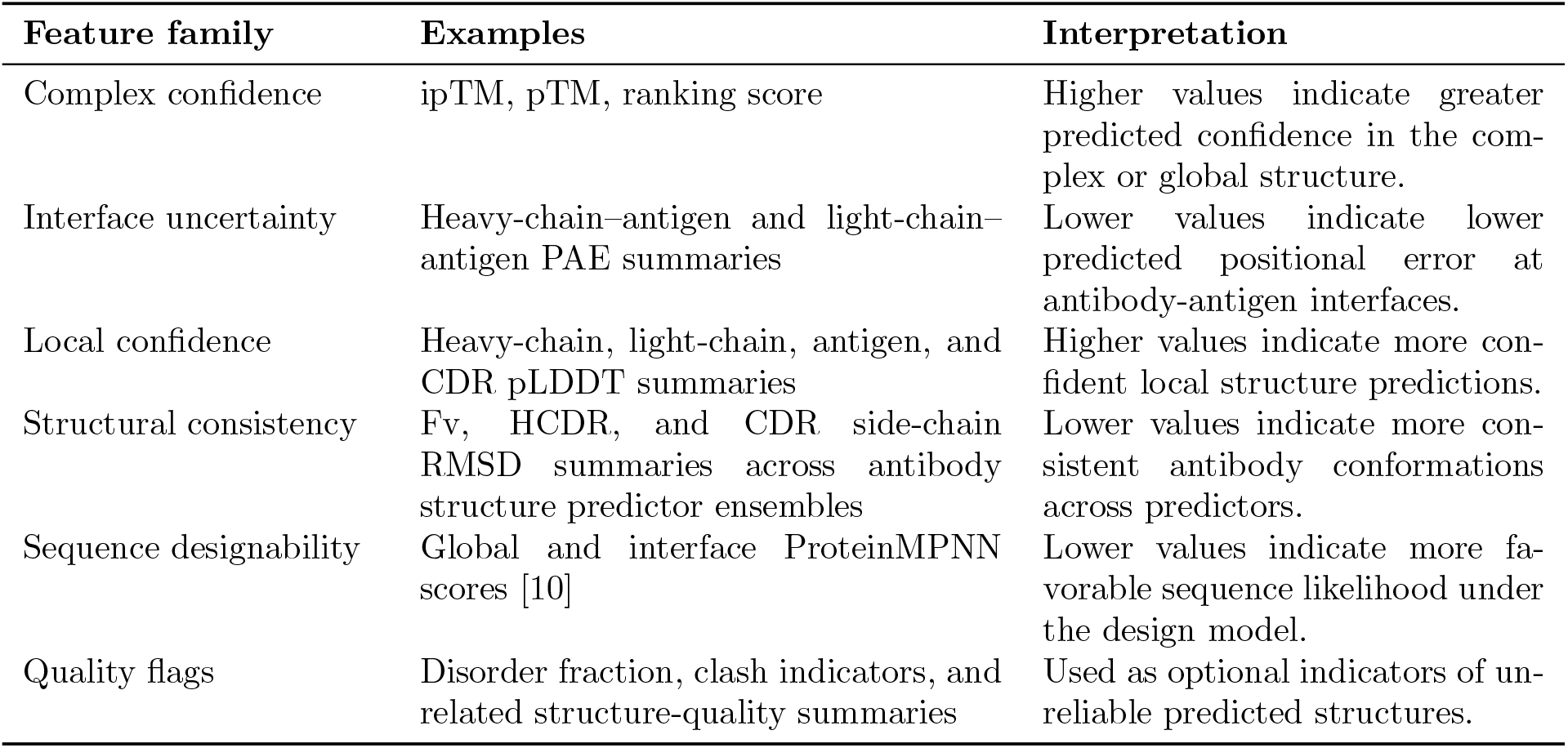
Feature families available to the rule-based filters.

| Feature family | Examples | Interpretation |
| --- | --- | --- |
| Complex confidence | ipTM, pTM, ranking score | Higher values indicate greater predicted confidence in the complex or global structure. |
| Interface uncertainty | Heavy-chain-antigen and light-chain-antigen PAE summaries | Lower values indicate lower predicted positional error at antibody-antigen interfaces. |
| Local confidence | Heavy-chain, light-chain, antigen, and CDR pLDDT summaries | Higher values indicate more confident local structure predictions. |
| Structural consistency | Fv, HCDR, and CDR side-chain RMSD summaries across antibody structure predictor ensembles | Lower values indicate more consistent antibody conformations across predictors. |
| Sequence designability | Global and interface ProteinMPNN scores [10] | Lower values indicate more favorable sequence likelihood under the design model. |
| Quality flags | Disorder fraction, clash indicators, and related structure-quality summaries | Used as optional indicators of unreliable predicted structures. |

**Table 3.** Performance comparison across methods, averaged over seven LOSO folds. ROC-AUC is the sole metric reported for this table.

| Category | Method | ROC-AUC | Training labels? |
| --- | --- | --- | --- |
| Rule-based | Exp00 baseline | 0.6371 | No |
| Rule-based | AF2-IG-easy | 0.6195 | No |
| Rule-based | Protenix-style adapted | 0.6612 | No |
| Rule-based | AF3-style adapted | 0.6471 | No |
| Rule-based | PXDesign-style | 0.5780 | No |
| Rule-based | **RMSD-Tuned Triad rule** | **0.8060** | **No** |
| ML | Logistic regression | 0.7144 | Yes |
| ML | Logistic regression + SelectKBest(50) + balanced weights | 0.7536 | Yes |
| LLM | GPT-4o full | 0.6508 | No |
| LLM | GPT-4o features | 0.5745 | No |
| LLM | GPT-4o tabular+few-shot | 0.7640 | No |
| LLM | GPT-5 tabular+few-shot | 0.8104 | No |

**Table 4.** Condensed milestones in the autoresearch trajectory.

| Experiment | Change | ROC-AUC | Gain vs. prior best |
| --- | --- | --- | --- |
| Exp00 | Initial hard-threshold baseline | 0.6371 | – |
| Exp01 | Adapted Protenix-style filter | 0.6612 | +0.0241 |
| Exp04 | Continuous soft scoring | 0.7280 | +0.0584 |
| Exp06 | Remove PAE; use ipTM, pTM, RMSD | 0.7605 | +0.0325 |
| Exp18 | Use minimum ensemble RMSD | 0.7805 | +0.0184 |
| Exp27 | Additive weighted power score, 3:1:3 | 0.7860 | +0.0003 |
| Exp53 | Multiplicative score | 0.7930 | +0.0020 |
| Exp62 | Increase ipTM emphasis | 0.7978 | +0.0015 |
| Exp66 | Tighten RMSD normalization to 1.3Å | 0.8008 | +0.0022 |
| Exp69 | Asymmetric exponents with RMSD power 0.35 | 0.8040 | +0.0015 |
| Final rule | Fine-tune RMSD power to 0.27 | **0.8060** | +0.0002 |

### 4.2 Autoresearch Progression

Figure 2 shows the expanded trajectory through Exp80, with the best result first reached by the RMSD-Tuned Triad rule. The largest qualitative change occurred when the search moved from binary hard thresholds to continuous scoring; later gains came from removing noisy features, using minimum ensemble RMSD instead of mean ensemble RMSD, replacing the additive weighted score with a multiplicative score, tightening the RMSD normalization scale, and then fine-tuning the RMSD exponent. The important point for this paper is not that a particular hand-written rule was obvious in advance, but that the loop made the search traceable: each retained improvement can be tied to a small design change and a fixed evaluation protocol.

**Figure 1.**
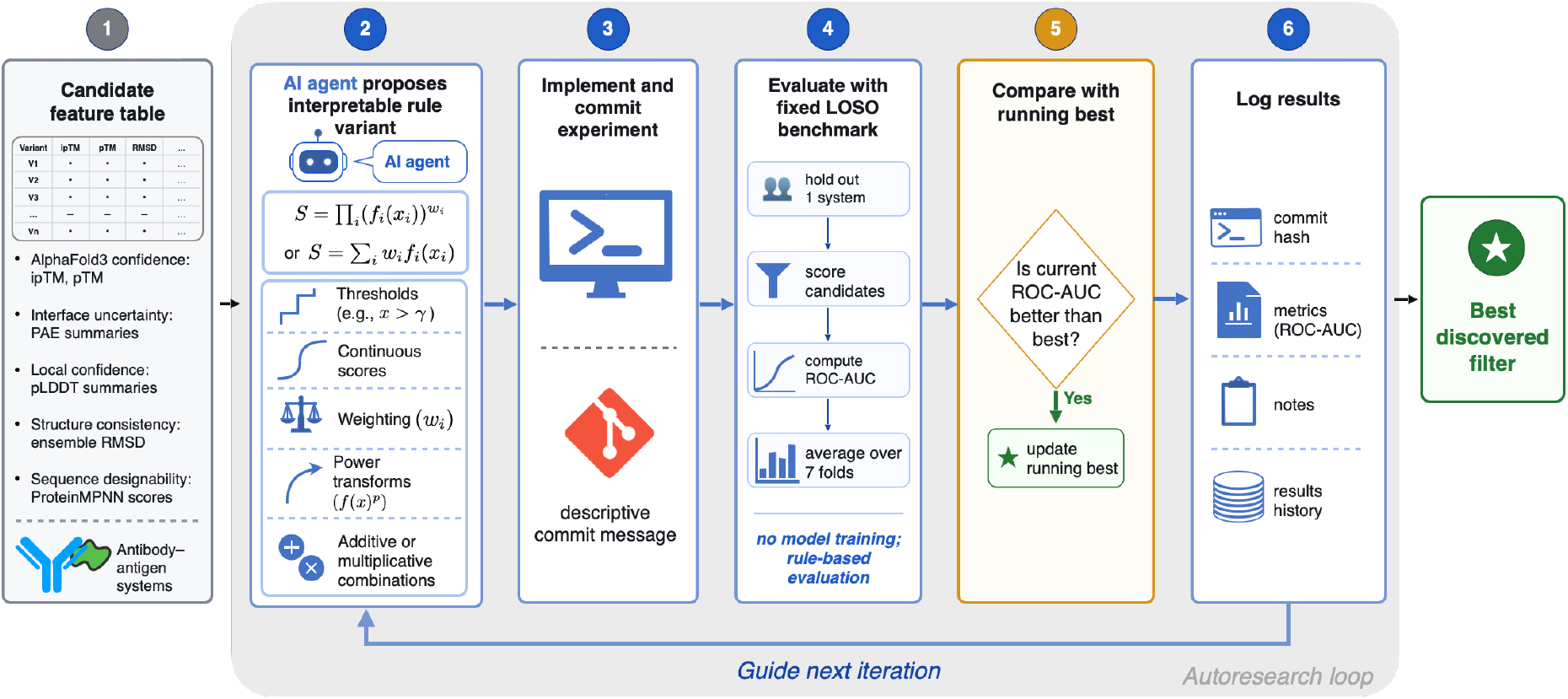
Autoresearch loop for rule discovery. Candidate-level structural and sequence scores define the available signals; each iteration proposes a rule variant, evaluates it under the fixed LOSO protocol, logs the result, and uses the running-best rule to guide the next proposal.

**Figure 2.**
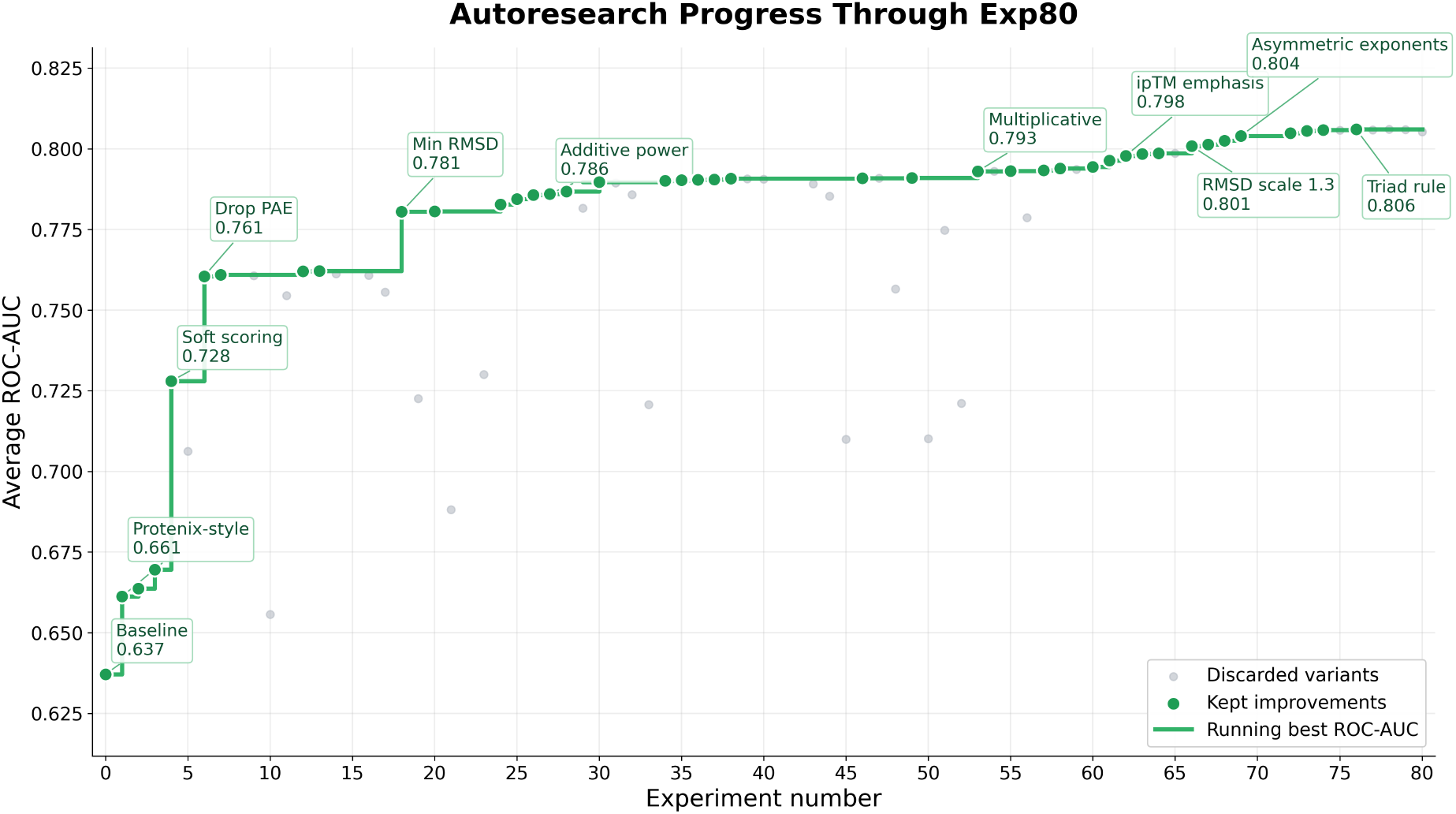
Autoresearch progress through Exp80. Green points indicate experiments that set a new best ROC-AUC, gray points indicate discarded variants, and the step line shows the running maximum. The search starts at 0.6371 ROC-AUC and reaches the final 0.8060 best with the RMSD-Tuned Triad rule.

### 4.3 Discovered Filter

The final filter is a compact multiplicative score over three normalized signals:

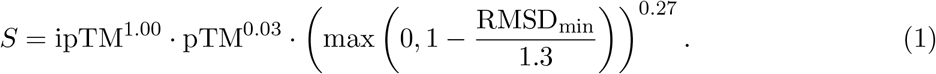

Here ipTM is the mean AF3 interface-confidence score for the antibody-antigen complex, pTM is the mean AF3 global structural-confidence score, and RMSD_min_ is the minimum Fv side-chain RMSD across the antibody structure predictor ensemble. These signals are defined in the appendix signal dictionary (Table 5). The RMSD term is therefore not an AF3 confidence metric; it measures structural consistency across antibody structure predictions. The ROC-AUC results use the continuous score directly, while a fixed operating threshold such as *S >* 0.5 can be chosen later if a binary screening decision is required. The score emphasizes interface confidence and structural consistency, with only a weak pTM exponent retained because removing pTM entirely reduced performance in a subsequent ablation check. Its main role in this paper is as the best artifact produced by the search loop.

**Table 5.**
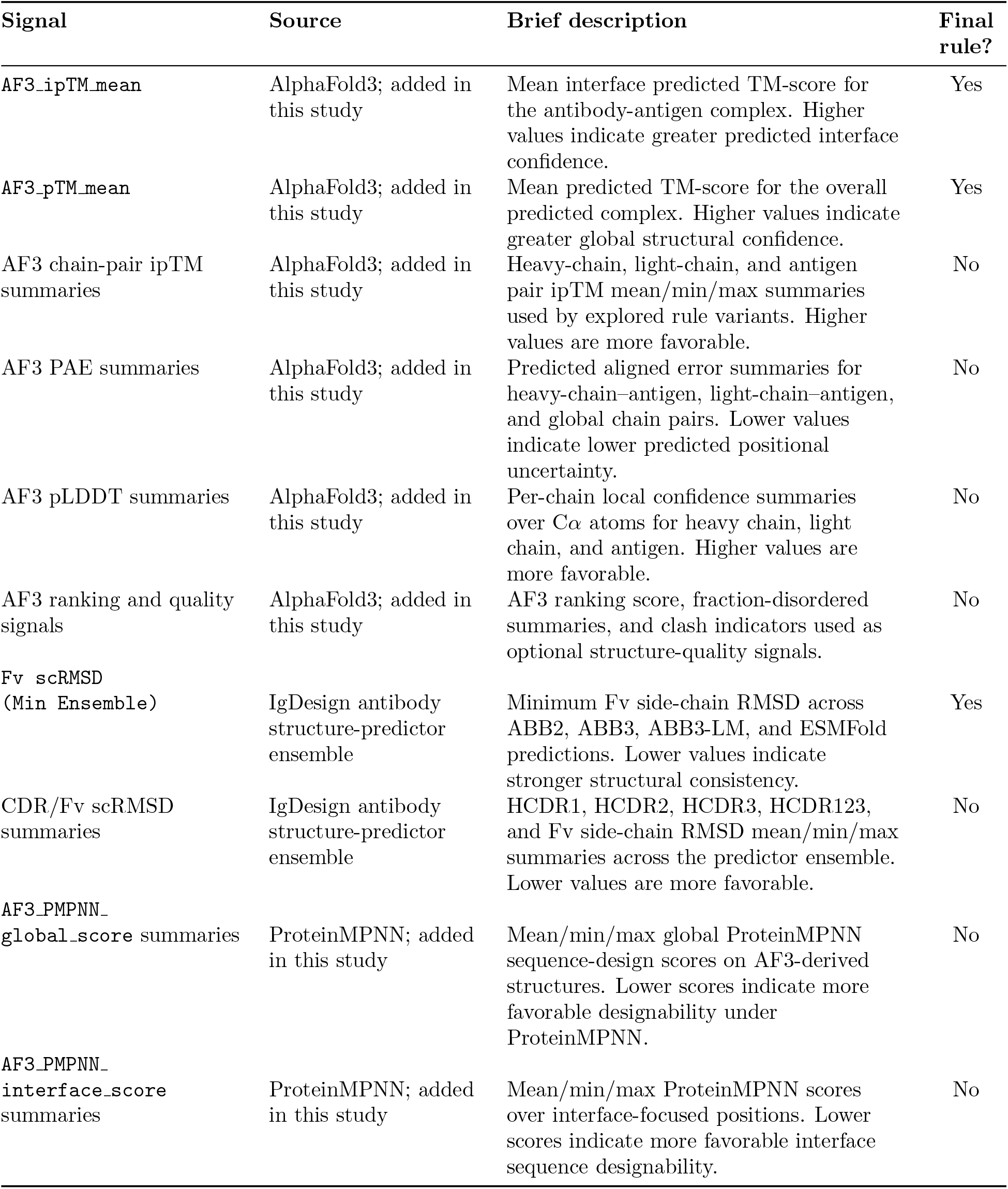
Signal dictionary for the rule-search feature set.

| Signal | Source | Brief description | Final rule? |
| --- | --- | --- | --- |
| AF3_ipTM_mean | AlphaFold3; added in this study | Mean interface predicted TM-score for the antibody-antigen complex. Higher values indicate greater predicted interface confidence. | Yes |
| AF3_pTM_mean | AlphaFold3; added in this study | Mean predicted TM-score for the overall predicted complex. Higher values indicate greater global structural confidence. | Yes |
| AF3 chain-pair ipTM summaries | AlphaFold3; added in this study | Heavy-chain, light-chain, and antigen pair ipTM mean/min/max summaries used by explored rule variants. Higher values are more favorable. | No |
| AF3 PAE summaries | AlphaFold3; added in this study | Predicted aligned error summaries for heavy-chain-antigen, light-chain-antigen, and global chain pairs. Lower values indicate lower predicted positional uncertainty. | No |
| AF3 pLDDT summaries | AlphaFold3; added in this study | Per-chain local confidence summaries over Cα atoms for heavy chain, light chain, and antigen. Higher values are more favorable. | No |
| AF3 ranking and quality signals | AlphaFold3; added in this study | AF3 ranking score, fraction-disordered summaries, and clash indicators used as optional structure-quality signals. | No |
| Fv_scRMSD (Min Ensemble) | IgDesign antibody structure-predictor ensemble | Minimum Fv side-chain RMSD across ABB2, ABB3, ABB3-LM, and ESMFold predictions. Lower values indicate stronger structural consistency. | Yes |
| CDR/Fv_scRMSD summaries | IgDesign antibody structure-predictor ensemble | HCDR1, HCDR2, HCDR3, HCDR123, and Fv side-chain RMSD mean/min/max summaries across the predictor ensemble. Lower values are more favorable. | No |
| AF3_PMPNN_global_score summaries | ProteinMPNN; added in this study | Mean/min/max global ProteinMPNN sequence-design scores on AF3-derived structures. Lower scores indicate more favorable designability under ProteinMPNN. | No |
| AF3_PMPNN_interface_score summaries | ProteinMPNN; added in this study | Mean/min/max ProteinMPNN scores over interface-focused positions. Lower scores indicate more favorable interface sequence designability. | No |

### 4.4 Mean ROC Curves

Figure 3 aggregates the RMSD-Tuned Triad rule, adapted rule-based baselines, and the GPT-5 tabular+few-shot LLM baseline across the seven held-out systems.

**Figure 3.**
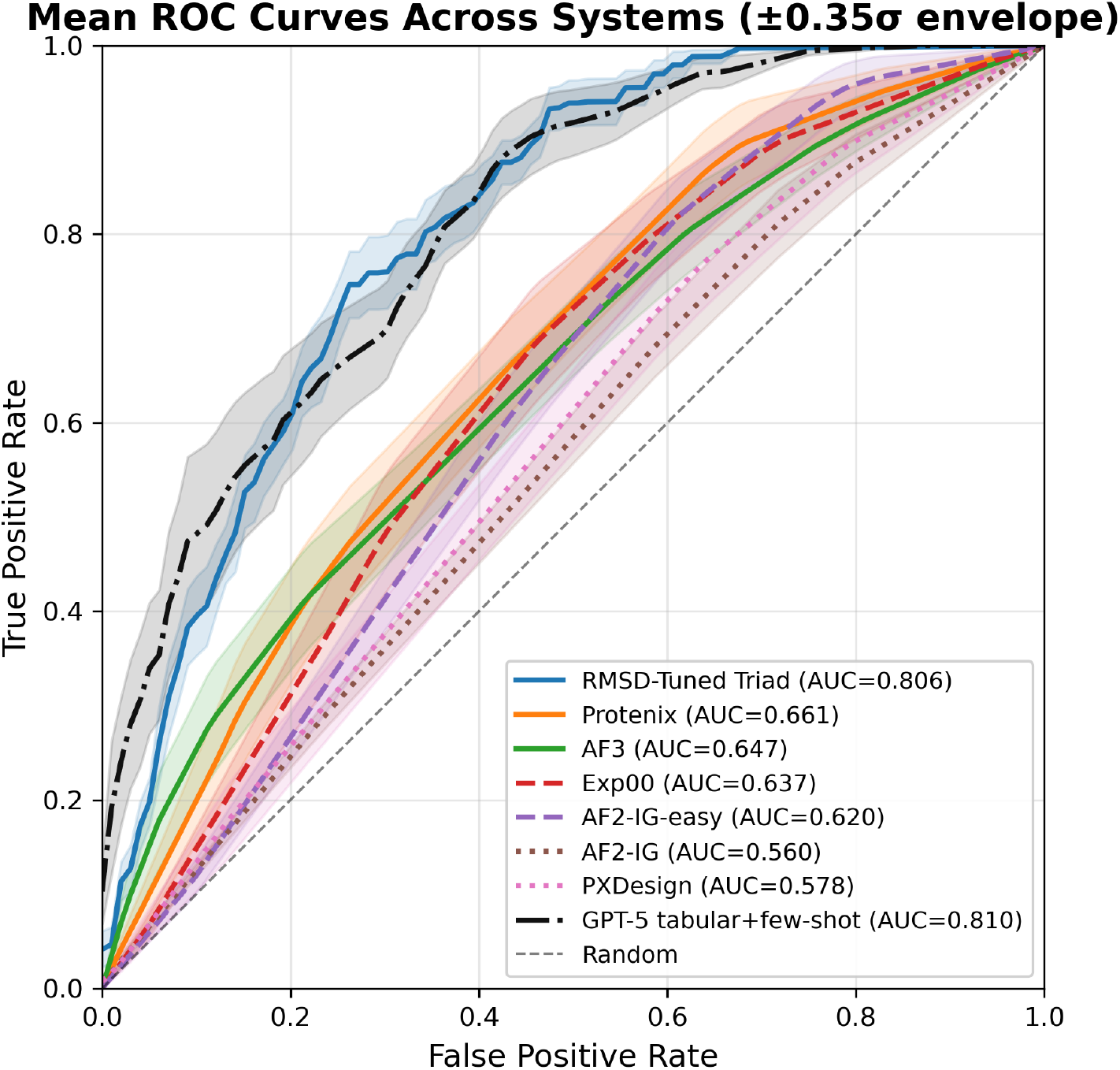
Mean ROC curves across systems with scaled standard-deviation envelopes. The RMSD-Tuned Triad curve shows the fixed discovered rule; the GPT-5 tabular+few-shot curve shows model-predicted binder probabilities. Adapted rule-based baselines are included for context.

**Figure 4.**
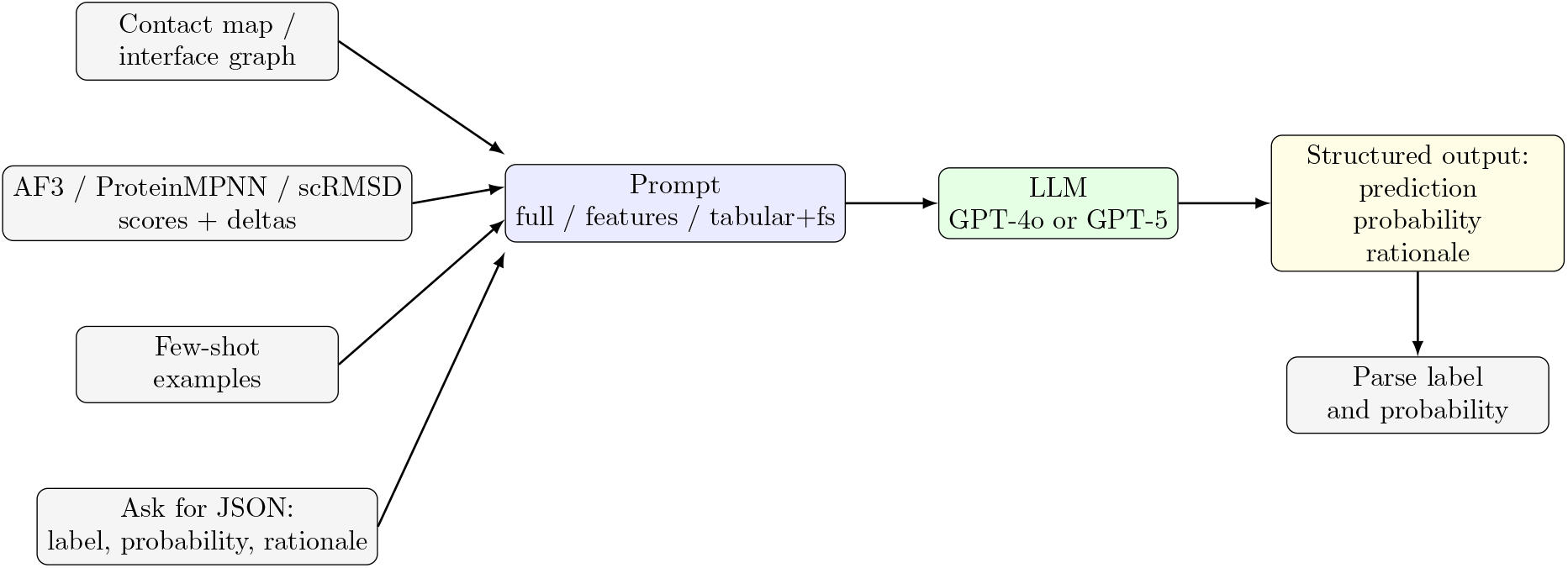
LLM binder-classification workflow. Interface structure, scalar AF3/ProteinMPNN/scRMSD features, and optional few-shot calibration examples are assembled into a prompt. The model returns a structured prediction whose probability field is used for ROC-AUC.

## 5 Discussion

The autoresearch loop worked because it preserved information that hard-threshold filters discarded. Binary AND rules treat a candidate barely above a cutoff the same as one far above it and reject candidates that fail any single metric by a small margin. Continuous scoring keeps the ordering information that ROC-AUC measures. The fractional power transformation further makes the filter less brittle by giving moderate confidence values useful credit, which matters when structure-prediction confidence is conservative for flexible antibody-antigen interfaces.

The comparison also clarifies the tradeoff between filter search, supervised learning, and LLM prompting. Supervised models can adapt to labeled systems, but in this benchmark the simple discovered filter still outperforms the two logistic-regression baselines. LLM prompting is flexible and can combine heterogeneous context, but it is more expensive, less deterministic, and sensitive to prompt format; the strongest GPT-5 prompt remains narrowly ahead by 0.0044 ROC-AUC, while the autoresearch filter gives essentially the same ranking quality with an explicit formula and no per-candidate LLM inference.

The study remains limited by the feature set and dataset size. The filter can only exploit signals already present in the AlphaFold3, structure-ensemble, and sequence-scoring outputs, and seven systems are not enough to claim universal transfer. The low threshold-dependent accuracy seen in earlier drafts is also a reminder that ROC-AUC and deployable decision thresholds are different questions. Future work should search operating thresholds for specific screening budgets, evaluate transfer to additional antibody design datasets, and test whether autoresearch-discovered scores are useful as features inside calibrated supervised models.

## 6 Conclusions

We presented an autoresearch loop for discovering interpretable rule-based filters for antibody binder classification. On seven IgDesign antibody-antigen systems, the loop improved average ROC-AUC from 0.6371 to 0.8060, a 26.5% relative gain over the initial baseline, without fitting to target-specific binding labels. The result is competitive with supervised ML and effectively matches the best LLM-prompting baseline while remaining compact, deterministic, and easy to inspect. This supports autoresearch as a practical way to improve filtering stages in antibody design workflows when labeled data are limited and explicit decision rules are preferred.

## Acknowledgments

This work was performed under the auspices of the U.S. Department of Energy by Lawrence Livermore National Laboratory under Contract DE-AC52-07NA27344 and was supported by the LLNL-LDRD Program under Project No. 24-ERD-022. LLNL-CONF-2015778.

## A Signal Dictionary

Table 5 summarizes the candidate-level signals exposed to the rule search. The final RMSD-Tuned Triad rule uses only three of these signals, but the autoresearch loop evaluated variants over the broader table.

## B Autoresearch Agent Prompt

The rule-search loop was driven by a repository-level prompt to Claude Code. This prompt was not used to classify individual antibody candidates. Instead, it defined the role of the coding agent that iteratively edited and evaluated the filter implementation. At a high level, the prompt specified four constraints: maximize mean LOSO ROC-AUC, remain within a rule-based filter search space, keep the evaluation protocol fixed, and record each experiment in version control and the results log.

The prompt described the seven IgDesign antibody-antigen systems, the available structure-derived and sequence-design signals, and the adapted literature-style baselines used as starting points. It emphasized that the search should use hard thresholds, soft thresholds, weighted scores, or related interpretable transformations, while explicitly prohibiting supervised machine learning models such as random forests, boosted trees, or neural networks.

Operationally, the prompt encoded the experimental loop shown in Figure 1: propose a small rule change, run the fixed LOSO evaluation, extract summary metrics from the run log, commit the implementation with a descriptive message, append the result to running text file, and use the observed outcome to choose the next rule variant. The prompt also included guardrails about metric directionality, normalization, threshold calibration, and overfitting to a single antibody-antigen system. These instructions made the search auditable: the final RMSD-Tuned Triad rule can be traced back through a sequence of explicit code changes and logged evaluations rather than being selected after an unrecorded manual sweep.

## C LLM Prompting Baseline Details

The LLM baselines use candidate-level prompts assembled from the same IgDesign and structure-derived evidence available to the rule-based filters, plus compact sequence and interface summaries. Each prompt is built from a system directory containing reference heavy- and light-chain variable-region sequences, the antigen sequence, a residue contact graph computed from the reference antibody-antigen structure, an interaction profile, optional sequence-scoring summaries, and AF3-extended candidate metrics. Antibody residues use IMGT-style labels in Chain+Position+AA format.

The contact graph block summarizes residue–residue contacts between antibody and antigen residues under a 7.0Åheavy-atom distance threshold. It reports chain IDs, total and per-chain edge counts, high-degree residues, and a truncated edge list with distances. The interaction-profile block summarizes hydrogen-bond, salt-bridge, and hydrophobic-contact counts. When sequence scoring is available, the prompt includes ProteinMPNN and language-model mutation-score summaries; lower minus-log-likelihood-style values are treated as more favorable sequence-design signals.

We evaluate four prompt/model variants. The **GPT-4o full** prompt includes raw reference sequences, candidate mutations, contact graph, interaction profile, sequence-scoring summaries, structure-ensemble summaries, and AF3 metrics. The **GPT-4o features** prompt removes most sequence prose and presents structured feature blocks, deltas, and heuristic cues. The **GPT-4o tabular+few-shot** prompt compresses the same evidence into numeric tables and prepends two binder and two non-binder examples from the same antigen system. The **GPT-5 tabular+fewshot** baseline uses the same tabular few-shot format with GPT-5. In all cases, the model is asked to return a JSON object containing a predicted label, a binder probability, and a short rationale; ROC-AUC is computed from the probability field.

## Prompt Excerpts

~~~
Reference antibody variable regions:
- Heavy: EVQLVESGGGLVQPG…WGRGTLVTVSS
- Light: EIVLTQSPATLSLS…FGGGTKVEIK
Contact graph:
chains heavy=H light=L antigen=G
total_edges=23 heavy_edges=13 light_edges=10
top_antibody_residues=H109G(4), H110G(3), …
edges:
H109G -- G127P (dist=6.563)
H110G -- G129I (dist=5.064)
Candidate changes vs reference:
  H105A->H105K; H106R->H106S; …; H117L->H117Y
Binding heuristics and cues:
 - CDR loops, especially CDRH3, are typical hotspots.
 - Aromatic residues in CDRs are enriched in paratopes.
 - Mutated contact hotspots: H109G(deg=4), H110G(deg=3)
Sequence scoring summary (minus_llr; lower is better):
 protbert: mean=0.01, min=-1.48, max=2.09
 abbert: mean=3.25, min=-0.78, max=7.13
TASK: classify binder vs non-binder; return JSON {prediction, probability, rationale}
SYSTEM: Afasevikumab-IL17A
CONTACTS: total_edges=23, heavy_edges=13, light_edges=10, threshold_A=7.0
HOTSPOTS (by degree): H109G=4, H110G=3, L108S=3, L109N=3, …
AF3 (mean/min/max):
 AF3_HC_ipTM_mean=0.480 (0.440..0.560)
 AF3_ipTM_mean=0.344 (0.240..0.500)
IGDESIGN scRMSD:
 ABB2_HCDR3=2.41, ABB2_Fv=0.84, …
~~~

